# The KIAA0319 Gene Polymorphisms are Associated with Developmental Dyslexia in Chinese Uyghur Children: Association of KIAA0319 Polymorphisms with Developmental Dyslexia

**DOI:** 10.1101/034660

**Authors:** Hua Zhao, Yun Chen, Bao-ping Zhang, Peng-xiang Zuo

## Abstract

To investigate the association of KIAA0319 gene polymorphisms and developmental dyslexia in individuals of Uyghurian descent. Eighteen single nucleotide polymorphisms (SNP) of gene KIAA0319 were screened in a group of 196 patients with dyslexia and 196 controls of Uyghur descent by determined the genotypes using a custom-by-design 48-Plex SNPscan™ Kit. SAS 9.1.3 software were used for data analysis. Seven SNPs(*P*_mm_=0.001) of KIAA0319 have significant differences between the cases and controls under specific genotype models. Especially for rs6935076(*P*_adjusted_=0.020 under dominant model; *P*_adjusted_=0.028 under additive model) and rs3756821(*P*_adjusted_=0.021 under additive model), which still associated with dyslexia after Bonferroni correction. The linkage disequilibrium analysis showed four block within gene KIAA0319 and only the ten-maker haplotype(*P=*0.013) in block 4 was significantly more common in dyslexia children than in controls. The results indicated that genetic polymorphisms of KIAA0319 are associated with increased risk of developmental dyslexia in Uyghur population.

## INTRODUCTION

Developmental dyslexia (DD) is a complex neuro-genetic disorder associated with impairment of reading performance despite adequate educational and intelligence opportunities, as well as in the absence of sensory or neurological disability. It is a common reading disability with the prevalence estimated to be 5%-17% of school-aged children in Western countries(Cope et al. 2005; Pennington and Bishop 2009). The corresponding figure for China, where the previous studies were carried out are reported to be 3.9%-8%(Sun et al. 2013). Although the etiology of dyslexia is not very clear, a number of studies shown that genetic factors play an important role in the development of dyslexia(Cope et al. 2005; Meng et al. 2005; Taipale et al. 2003; Hannula-Jouppi et al. 2005). So far, there are more than nine dyslexia susceptibility loci have been mapped and allocated based on linkage studies (From DYX1 to DYX9). Subsequent association studies have identified several candidate genes at most of these locus, including KIAA0319(Cope et al. 2005), DCDC2(Meng et al. 2005), DYX1C1(Taipale et al. 2003) and ROBO1(Hannula-Jouppi et al. 2005).

KIAA0319, located at chromosomes 6p22.2–22.3. It is first proposed by Kaplan et al. (Kaplan et al. 2002), who found a microsatellite marker residing in KIAA0319 according to linkage studies. Then, dyslexia-KIAA0319 association studies are conducted in US(Francks et al. 2004), UK(Harold et al. 2006; Cope et al. 2005), German(Ludwig et al. 2008), Canadian(Couto et al. 2010; Elbert et al. 2011), Indian(Venkatesh et al. 2011; Venkatesh et al. 2013) and Chinese(Lim et al. 2014; Sun et al. 2014) population, suggesting several risk SNPs are associated with reading disability, such as rs4504469, rs6935076 and rs2038137. However, the results are not always consistent since the difference of genetic and linguistic between ethnic groups. KIAA0319 gene is expressed in human cerebral neocortex(Paracchini et al. 2006). The protein encoded by KIAA0319 has a large, highly N-and O-glycosylated plasma membrance and is involved in abnormal regulation of neuronal migration and neurite growth through endocytosis pathway of the protein and proteolytic processing(Velayos-Baeza et al. 2008; Levecque et al. 2009; Velayos-Baeza et al. 2010), which is considered to be an important feature of dyslexia(Gabel et al. 2010; Poelmans et al. 2011). However, both the exact pathogenesis of candidate gene KIAA0319 and the mechanism of dyslexia associated KIAA0319 polymorphisms are remains complicated.

Thus far, most studies on dyslexia associated polymorphisms have been performed in European and Chinese populations and few covers individuals of Uyghurian descent. The Uyghur, accounts for 48% of the whole group in Xinjiang with 11 million population, is the second largest minority ethnic group in China, mainly live in Xinjiang Uyghur Autonomous region, areas are located in far northwest of China(Statistic Bureau of Xinjiang Uyghur Autonomous 2014). It is a population presenting a typical mixed genetic origin, both Eastern and Western anthropometric traits (Black et al. 2006; Wang et al. 2003). Uighurs have their own inheritance, culture, religion and language, which are very different from other ethnic populations. Based on our previous epidemiology and genetic studies, we performed case-control association study among a large unrelated Chinese Uyghur cohort in the present research, to investigate whether there is also an association of KIAA0319 gene polymorphisms and Uyghur dyslexia children.

## MATERIALS AND METHODS

### Experimental subjects

We selected 4251 Uyghur primary students aged 8–13 years by cluster sampling in Kashgar and Aksu, Xinjiang, China. The study was approved by the ethical committee of The Medical School of Shihezi University. Informed written consents were obtained from all participants students and their guardians. The diagnostic criteria for dyslexia were based on the following criteria: ① the score of *The Pupil Rating Scale Revised Screening for Learning Disability* was lower than 65 points(Jing et al. 1998); ②a score on *The Dyslexia Checklist for Uyghur Children* at least two standard deviations higher than the mean score(Wu et al. 2006); ③an intelligence quotient score was higher than 80 assessed by *China-Wechsler Intelligence Scale* for Children(Gong and Dai 1987); and ④in the absence of visual and/or auditory disorders or psychiatric diseases. In total, 228 Uyghur students were diagnosed as dyslexic and 196 of which were participant in the present study, with a total 122 boys and 74 girls, aged between 8 and 12 years(mean age=10.99±1.1 years). Then age-, education-, gender- and ethnicity-matched 196 normal children were also recruited for the case-control study.

### Genotyping

A total of eighteen SNPs of KIAA0319 were selected in the present study for the following reasons. ①The minor allele frequencies (MAF) of these eighteen SNPs were more than 0.05 in both HapMap CHB data and HapMap CEU data. ②These polymorphisms were reported to be associated with dyslexia in previous study among Indo-European(rs4504469; rs2038137; rs6935076; rs2179515; rs3212236; rs761100; rs9461045)(Cope et al. 2005; Harold et al. 2006; Couto et al. 2010; Venkatesh et al. 2013) and Chinese language(rs1091031; rs699463; rs3903801; rs12193738; rs2760157; rs807507; rs16889506; rs9366577; rs16889556; rs2038139; rs3756821)(Sun et al. 2014; Lim et al. 2014).

DNA was extracted from oral mucosal cells by the buccal swabs method as described elsewhere(Zuo et al. 2012). The SNP genotyping work was performed using a custom-by-design 48-Plex SNPscanTM Kit(Cat#: G0104K, Genesky Inc. Shanghai, China) as what mentioned before(Chen et al. 2012). This kit was developed according to patented SNP genotyping technology which is based on double ligation and multiplex fluorescence PCR. Our study data were collected through procedures carried out according to the manufacturer’s manual. Each 96-well plate included one non-template control. For quality control, repeated analyses were done for 4% of randomly selected samples with high DNA quality. Call rates for each SNP were above 98%.

### Statistical analysis

The Hardy–Weinberg equilibrium(HWE) tests were performed for each SNP. Differences in the distribution of demographic characteristics, selected variables, and genotypes of the 18 SNPsbetween the cases and controls were evaluated using the χ^2^ test. The associations analysis between SNPs genotypes and risk of dyslexia were estimated by computing the ORs and their 95% CIs using logistic regression analyses with dominant, recessive, over-dominant, additive, and genotype models. Besides, Linkage disequilibrium(LD) analysis of eighteen SNPs and haplotype selection were performed using Haploview software (Version 4.2)(Barrett et al. 2005). Bonferroni correction was applied for multiple comparisons. All statistical analyses were performed with SAS 9.1.3 (SAS Institute, Cary, NC, USA) and P values were twotailed with a significance level of 0.05.

## RESULTS

### Characteristics of the Uyghur population

On the whole, 195 cases and 196 controls were successfully genotyped in the study with a response rate is 99.5% and 100%, respectively. Characteristics of these Uyghur students were summarized in Table 1. The dyslexia and normal students appeared to be adequately matched on age, sex, education and ethnic as suggested by the χ^2^ tests (*P*>0.05). Besides, the primary information of the eighteen genotyped SNPs is shown in Table 2. The MAF of most SNPs in control students was between the MAF of CHB data and CEU data. All the markers showed HWE *P*-value>0.05, except rs16889506(*P=*0.022).

**Table 1.** Distribution of selected demographic variables and risk factors of participants

| Variable | Cases(n=195) | | Control(n=196) | | $\chi^2$ | P |
| --- | --- | --- | --- | --- | --- | --- |
|  | n | % | n | % |  |  |
| Age |  |  |  |  | 0.124 | 0.725 |
| $\leq 10$years | 76 | 38.97 | 73 | 37.24 | | |
| $\geq 11$years | 119 | 61.03 | 123 | 62.76 | | |
| Sex |  |  |  |  | 0.346 | 0.556 |
| Boys | 125 | 64.10 | 120 | 61.22 |  |  |
| Girls | 70 | 35.90 | 76 | 38.78 |  |  |
| Grade |  |  |  |  | 0.005 | 0.998 |
| Three | 56 | 28.72 | 56 | 28.57 |  |  |
| Four | 69 | 35.38 | 70 | 35.71 |  |  |
| Five | 70 | 35.90 | 70 | 35.71 |  |  |

**Table 2.** List of SNPs and HWE’s of *KIAA0319* analyzed by SNPscan in the present study

| SNP | Regulome DB | Allele | Position | Location | CHB/CEU<sup>b</sup> | Control | HWE<sup>d</sup> |
| --- | --- | --- | --- | --- | --- | --- | --- |

|  | score<sup>a</sup> |  |  |  | MAF<sup>c</sup> | MAF |  |
| --- | --- | --- | --- | --- | --- | --- | --- |
| rs1091031 | No Data | G/A | 24539139 | 3 'downstream | 0.193/0.438 | 0.298 | 1.000 |
| rs699463 | 4 | G/A | 24544903 | Exon21 | 0.121/0.239 | 0.184 | 1.000 |
| rs3903801 | 5 | A/G | 24559433 | Intron16 | 0.246/0.429 | 0.327 | 0.330 |
| rs12193738 | No Data | T/C | 24568393 | Intron13 | 0.215/0.473 | 0.324 | 0.145 |
| rs2760157 | 5 | G/A | 24578272 | Exon 9 | 0.489/0.183 | 0.332 | 0.262 |
| rs807507 | No Data | C/G | 24579867 | Intron 8 | 0.178/0.483 | 0.309 | 0.093 |
| rs4504469 | No Data | C/T | 24588884 | Exon 4 | 0.019/0.306 | 0.250 | 0.339 |
| rs16889506 | 1f | T/C | 24595853 | Intron 3 | 0.109/0.186 | 0.219 | 0.022 |
| rs2179515 | No Data | C/T | 24628203 | Intron 1 | 0.113/0.385 | 0.265 | 0.714 |
| rs761100 | 6 | C/A | 24632642 | Intron 1 | 0.130/0.455 | 0.304 | 0.502 |
| rs9366577 | No Data | T/C | 24641328 | Intron 1 | 0.073/0.150 | 0.082 | 0.120 |
| rs16889556 | No Data | C/T | 24641605 | Intron 1 | 0.201/0.155 | 0.122 | 1.000 |
| rs6935076 | 5 | C/T | 24644322 | Intron 1 | 0.230/0.262 | 0.212 | 0.668 |
| rs2038139 | No Data | A/C | 24645420 | Intron 1 | 0.122/0.332 | 0.265 | 1.000 |
| rs2038137 | 2b | G/T | 24645943 | 5' UTR | 0.234/0.334 | 0.265 | 1.000 |
| rs3756821 | 4 | C/T | 24646821 | 5' UTR | 0.411/0.362 | 0.276 | 0.593 |
| rs3212236 | 5 | T/C | 24648455 | Promoter | 0.358/0.358 | 0.452 | 0.774 |
| rs9461045 | No Data | C/T | 24649061 | Promoter | 0.481/0.221 | 0.452 | 0.774 |

a:http://www.regulomedb.org/.
b: CHB/CEU: Han Chinese in Beijing, China/ Utah residents with ancestry from northern and western Europe
c:MAF: minor allele frequency.
d:HWE: Hardy–Weinberg equilibrium.

### Associations between polymorphisms and risk of dyslexia

In the present study, seven of the eighteen KIAA0319 polymorphisms showed nominal association with dyslexia after genotyping (results shown in Table 3). Allelic frequencies of six SNPs(rs1091031*:P=*0.034; rs16889556*:P=*0.010; rs6935076*:P=*0.001; rs3756821*:P=*0.001; rs3212236*:P=*0.009; rs9461045*:P=*0.009) were significant different between dyslexia and control students. Furthermore, the minor allele(T) frequency of rs6935076 with *P=*0.026 and T allele of rs3756821 with *P=*0.023 also displayed a strong association with dyslexia after applying Bonferroni’s correction.

Table 3 also shows genotype distributions under various genotype models of these seven risk SNPs. The results showed significant association of rs1091031 in dominant model(*P=*0.042) and additive model(*P=*0.033). SNP rs9366577 demonstrated nominal significant association under over-dominant model(*P=*0.024). SNP rs16889556 showed significant association under dominant(*P=*0.007), over-dominant(*P=*0.012), additive model(*P=*0.009) and in heterozygous genotype of co-dominant model(*P=*0.010). Similarly, SNP rs6935076 manifested nominal meaning association based on dominant(*P=*0.001), over-dominant(*P=*0.009) and additive model(*P=*0.001) as well as in co-dominant model(CT vs CC: *P=*0.003; TT vs CC: *P=*0.046). Besides, significant association was also found for rs3756821 under dominant(*P=*0.004), recessive(*P=*0.021), additive model(*P=*0.001) and co-dominant model(CT vs CC: *P=*0.021; TT vs CC: *P=*0.004). Moreover, SNP rs3212236 and rs9461045 showed equal significant association under dominant(*P=*0.019) and additive model(*P=*0.010), as well as in homozygous genotype of co-dominant model(*P=*0.015). However, after Bonferroni correction for multiple comparisons in different models, only polymorphism rs6935076 under dominant model(*P=*0.020) and additive model(*P=*0.028) as well as rs3756821 under additive model(*P=*0.021) remain significant difference between dyslexia and controls.

**Table 3.**
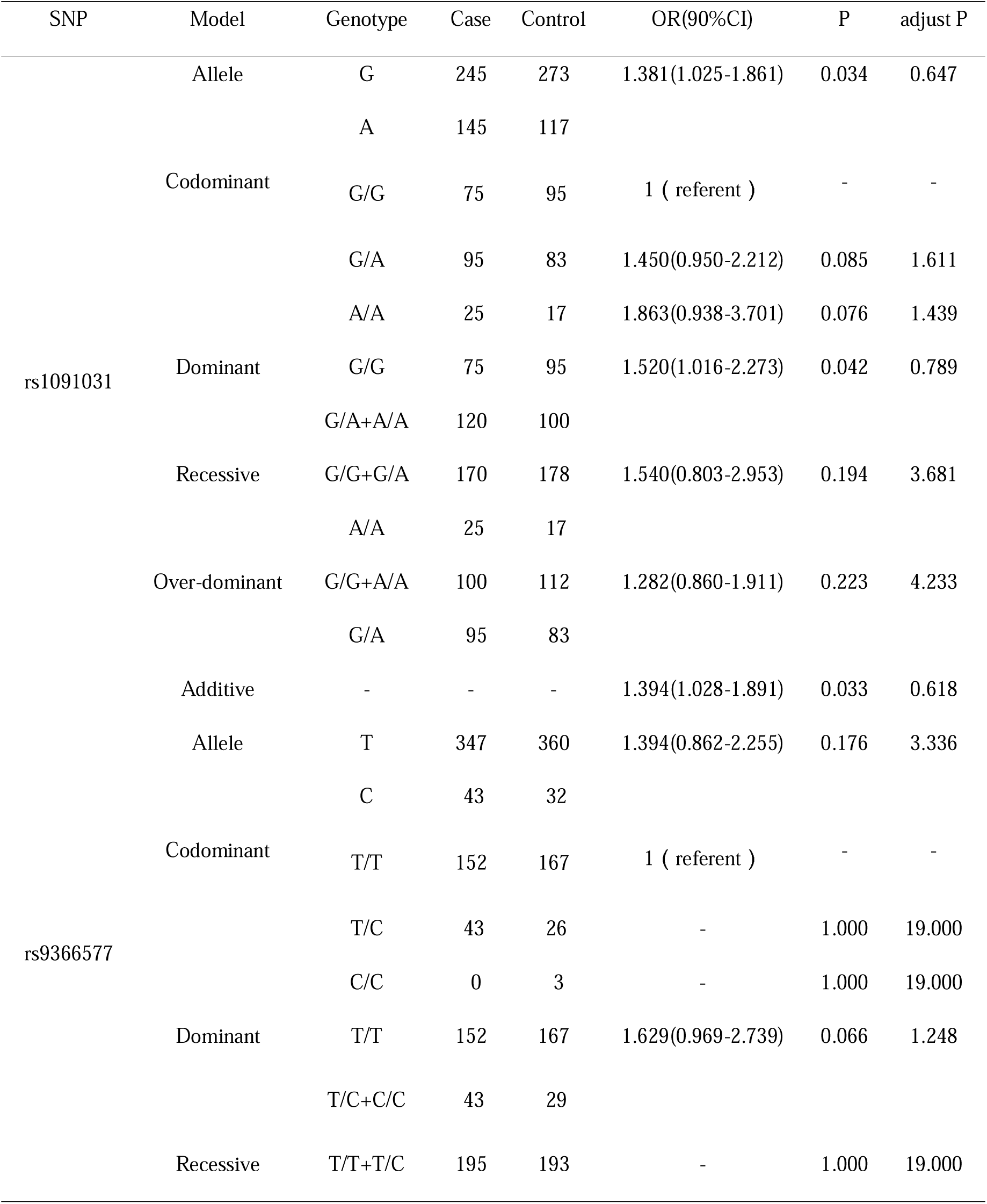
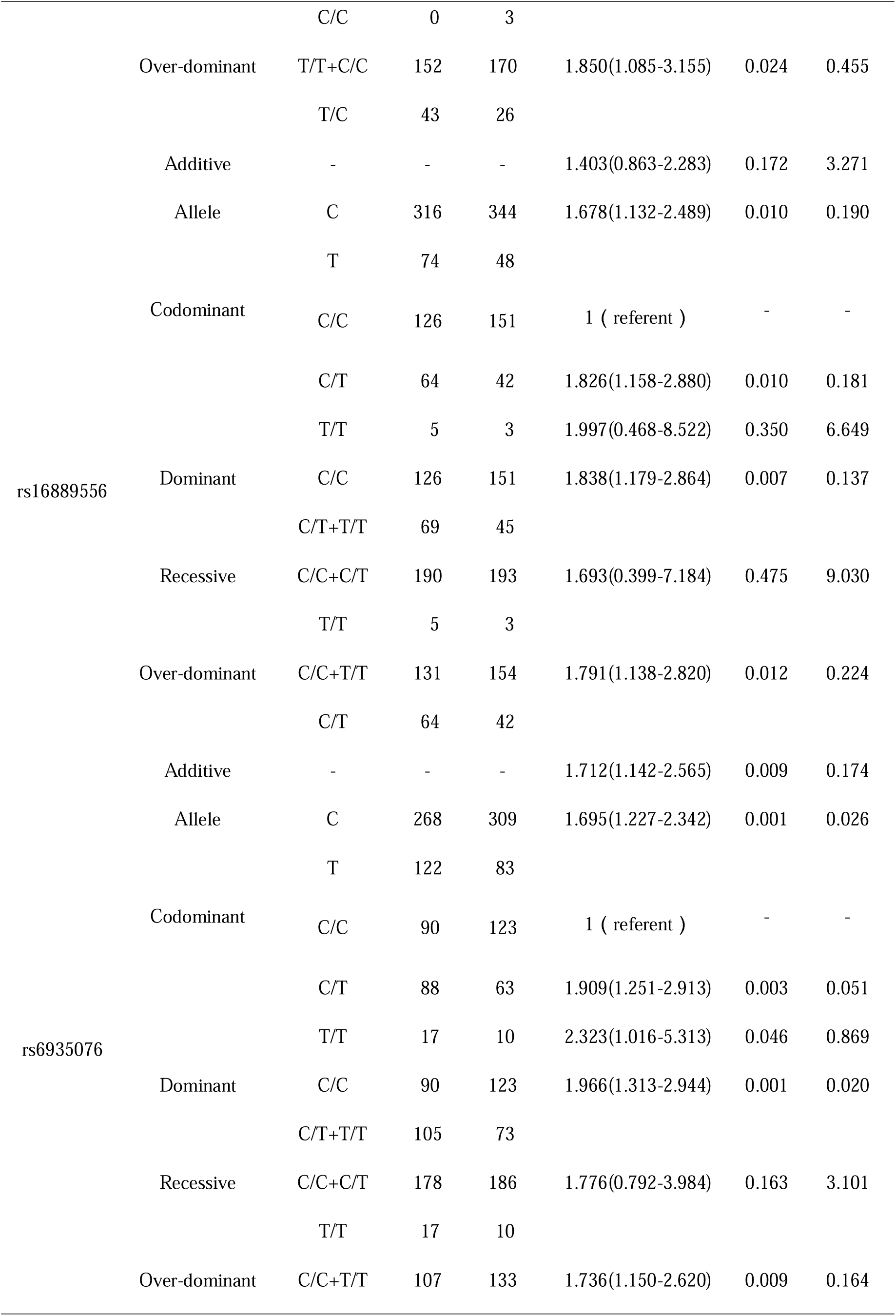
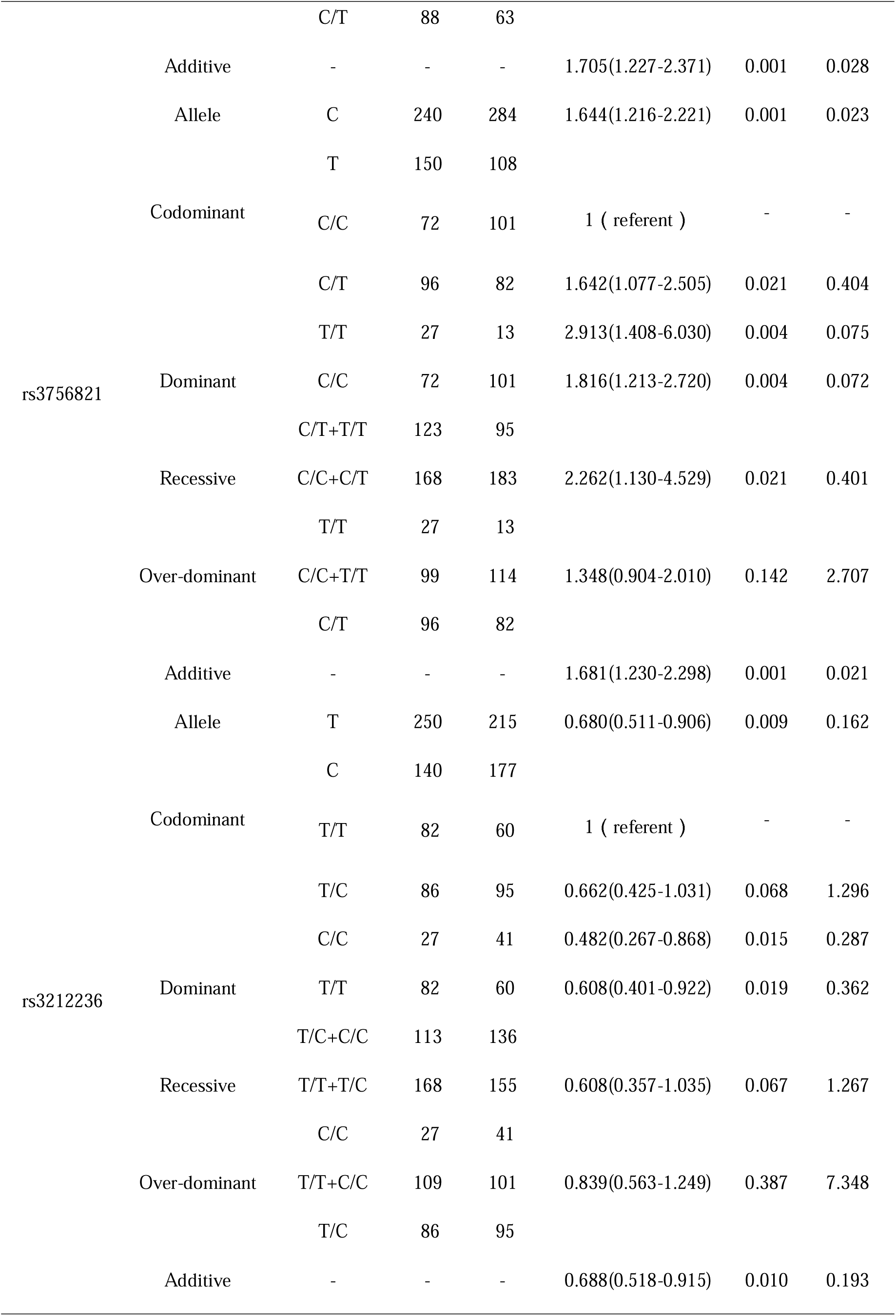
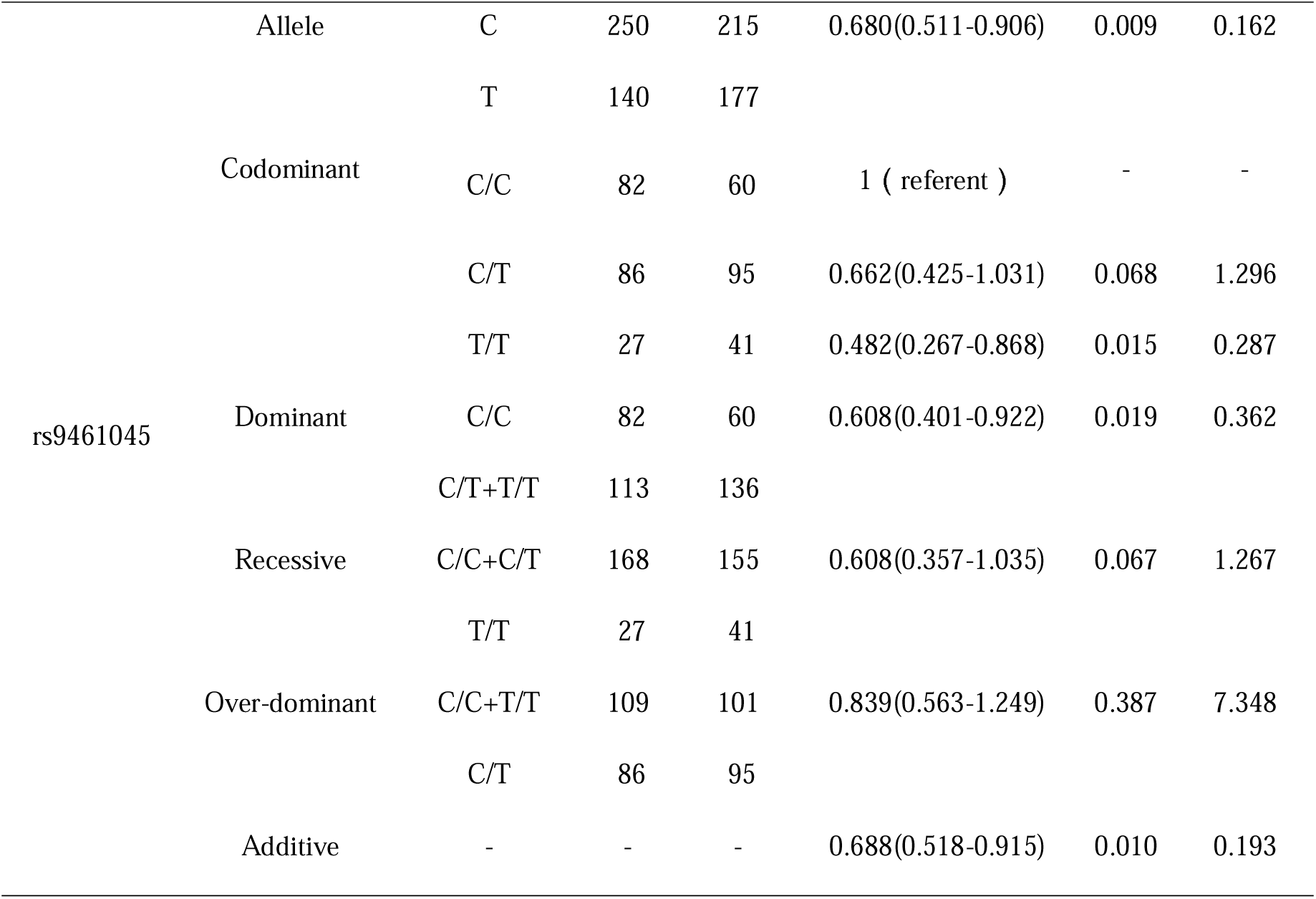
Logistic regression analyses of association between KIAA0319 polymorphisms and risk of dyslexia

### Haplotype analysis

Four detected LD blocks within gene KIAA0319 are shown in Fig. 1 according to D’ value. Table 4 indicates the frequency of the CCTTTAGTTC haplotype of block 4 in dyslexia group was significantly higher than that in control group(*P=*0.013) and implies that this haplotype was a risk haplotype to dyslexia. Except for block 4, we did not find any significant haplotypes in all the other three blocks.

**Table 4.** Selected haplotype analysis results of KIAA0319

| Haplotype | Case | Control | OR (95%CI) | P-value |
| --- | --- | --- | --- | --- |
| Block 1: rs699463-rs3903801 |  |  |  |  |
| GA | 190 | 192 | - | - |
| GG | 114 | 128 | 0.9000(0.6520-1.2424) | 0.5218 |
| AA | 86 | 72 | 1.2070(0.8322-1.7506) | 0.3213 |
| Block 2: rs12193738-rs2760157-rs807507 |  |  |  |  |
| TGC | 159 | 136 | - | - |
| TAC | 110 | 129 | 0.7294(0.5180-1.0270) | 0.2364 |
| CGG | 115 | 121 | 0.8129(0.5770-1.1454) | 0.0707 |
| Block 3: rs4504469-rs16889506 |  |  |  |  |
| CT | 195 | 208 | - | - |
| TT | 95 | 98 | 1.0340(0.7336-1.4574) | 0.8485 |
| CC | 100 | 86 | 1.2403(0.8756-1.7569) | 0.2254 |
| Block 4: rs2179515-rs761100-rs9366577-rs16889556-rs6935076-rs2038139-rs2038137-rs3756821-rs3212236-rs9461045 |  |  |  |  |
| TATCCCTCTC | 89 | 103 | - | - |
| CCTCCAGCCT | 140 | 176 | 1.0597(0.6955-1.6145) | 0.7872 |
| CCTTTAGTTC | 73 | 47 | 1.7975(1.1308-2.8573) | 0.0131 |

**Figure1.**
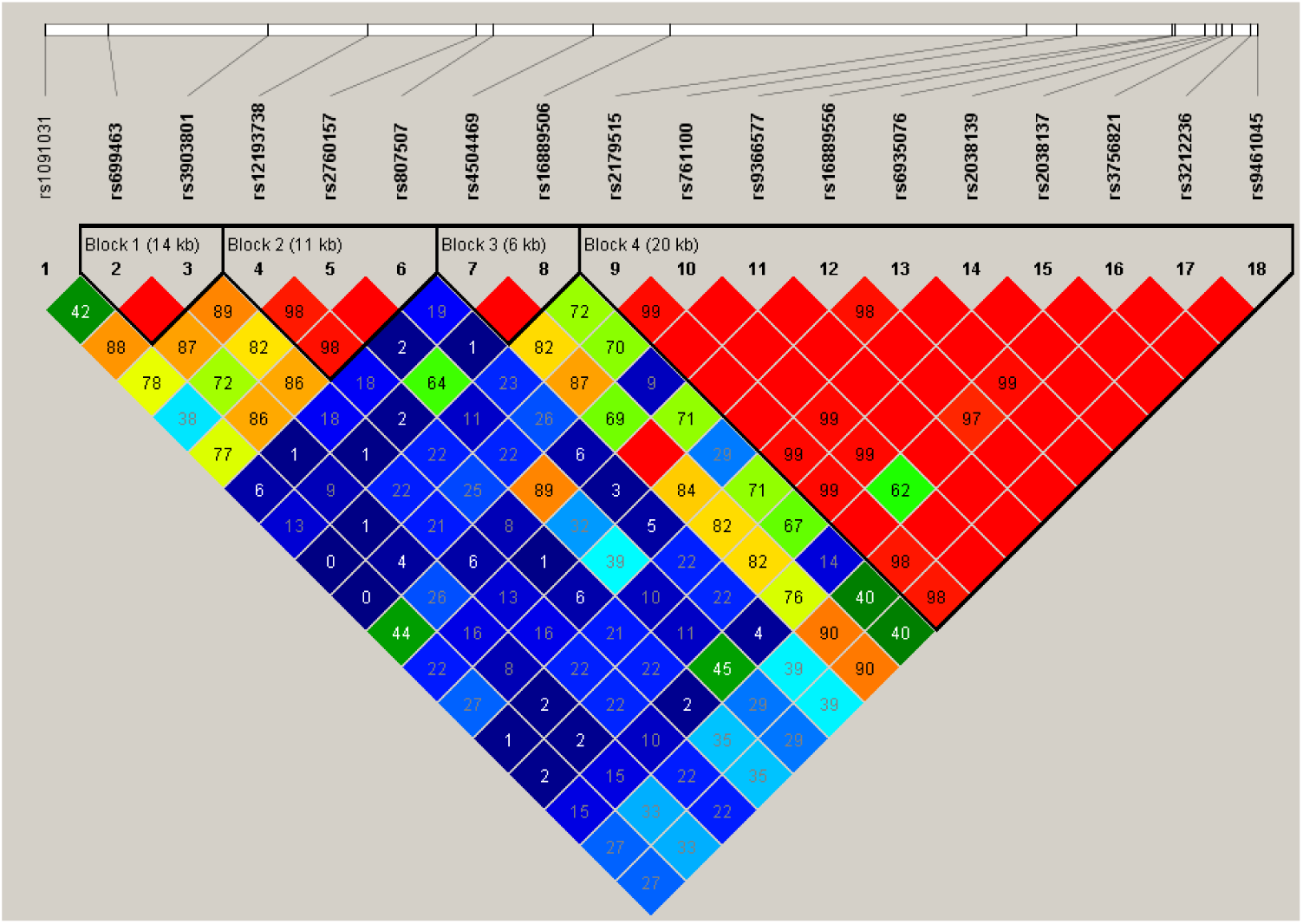
Linkage disequilibrium (LD) block generated by the Haploview 4.1. Regions of low-to-high LD, as measured by the D statistic, were represented by deep blue to red shading, respectively. LD blocks were analyzed using an algorithm by Gabriel et al.(Gabriel et al. 2002)

## DISCUSSION

Up to now, there are numerous studies focused on dyslexia associated KIAA0319 polymorphisms. However, these studies mainly performed in European, Indian and Han Chinese populations and the results are inconclusive. Take the genetic and linguistic differences between Uyghur and other populations into consideration, here we selected eighteen tag SNPs of gene KIAA0319 followed by high-throughput sequencing to assess the association of polymorphisms and dyslexia in a large unrelated Uyghur cohort. In summary, we identified seven KIAA0319 polymorphisms and one haplotype have nominal association with dyslexia after genotyping. Especially for rs3756821 and rs6935076, the results survived Bonferroni correction for multiple comparisons. To our knowledge, this study is first to explore the role of KIAA0319 in dyslexia in a relatively large Uyghur populations.

This study included frequently reported polymorphisms by previous study among Chinese and Indo-European language. The rs3756821, located at 5’ UTR of gene KIAA0319, showed nominal association with dyslexia in recent studies in Chinese populations but failed showed significant association after Bonferroni correction(Lim et al. 2014; Sun et al. 2014). While the results from the present study indicate the association between rs3756821 and Uyghur dyslexia after correction. Therefore, it implies that rs3756821 is a risk SNP to dyslexia in Uyghur population not in Chinese. This may be attributed to the fact that the genetic and linguistic among Uyghur and Chinese might be different, which could cause the susceptibility gene conduce to dyslexia through different mechanisms(Kirsten et al. 2012). Compared to Chinese, Uyghur form an isolated genetic group because of the Uyghur population is less influenced by recent migration(Black et al. 2006) and Uyghur are overwhelmingly Muslim, who prohibits marriage to non-Muslims(Li 2012). For language, Uyghur, an alphabetical language, belongs to the Altaic family with linear one-dimensionally arranged alphabetic(Xi et al. 2015), which is dislike Chinese (ideograph language) in linguistic characteristics(Siok et al. 2008). Besides, previous studies found that the cerebral regions activated by Uyghur and Chinese language are not identical and identified the left anterior cinugulate gyrus might associated with Uyghur language(Xi et al. 2015).

We also find significant association of rs6935076, a previously identified causative polymorphism with dyslexia. rs6935076 is located in intron 1 of the gene KIAA0319 and has the role to explain dyslexia status in the research done by Cope et al. among UK populations(Cope et al. 2005). However, several previous association study(Brkanac et al. 2007) and our early meta analysis failed to replicate the association of rs6935076 and dyslexia. Except for rs3756821 and rs6935076, other selected polymorphisms seem unlikely contributing to dyslexia susceptibility among Uyghur children after Bonferroni correction, whereas the results are inconclusive across different populations(Venkatesh et al. 2013; Elbert et al. 2011; Sun et al. 2014). The genetic and linguistic also could account for this results. We have mentioned that Uyghur presenting a typical mixed genetic origin and is an isolated genetic group. Uyghur and English both are part of an alphabetical language, but Uyghur has its own written forms and rules for writing and it is more complex than English(Siok et al. 2008). Even our findings reveal that rs3756821 and rs6935076 have significant difference between dyslexia and control group, the mechanism of these variants in Uyghur dyslexics is still unclear.

It is generally believed that combined variants within a gene may provide a more comprehensive evaluation than a single polymorphism in association studies(Pan et al. 2010). Based on the analysis of haplotypic, we think that the susceptibility haplotype (CCTTTAGTTC) constructed by ten SNP has an independent effect on dyslexia(P*=*0.013). However, we should take care of the results to assess the association between the haplotype and dyslexia because Fallin et al.(Fallin and Schork 2000) suggested that deviation from HWE with a high level of heterozygosity may produce error genetic effect of haplotype. Therefore, we concluded that children who carry more risk alleles are more likely to develop dyslexia than those who carry fewer or none.

While the present study provides valuable insight into the genetic differences of dyslexia in minority group, several limitations need to be addressed. First of all, the SNPs selection were based on previous study, we devoted ourselves to verified whether these polymorphisms have the same effect on Uyghur dyslexia children, which may not give a comprehensive view about genetic variability of KIAA0319 in Uyghur population. Secondly, the moderate sample size could limited the statistic power of this study. There were actually more dyslexia children included in the present study, but only 86% of dyslexia children were participate in the study due to the people of minority group often refuse to join in scrape the oral mucosal. Moreover, dyslexia is a multiple etiology disease, the existence of the interaction between environmental factors and the dyslexia susceptibility loci, as well as the interaction between different candidate genes must be further validated in the Uyghur population.

In conclusion, we performed association study of KIAA0319 with dyslexia in a large unrelated Uyghur cohort through SNPs selection and genotyping. Our study suggested that seven polymorphisms and one haplotype of KIAA0319 have significant association with the risk of dyslexia. Especially for rs6935076 and rs3756821, which still associated with dyslexia after Bonferroni correction. Results could contribute to early identification and management of Uyghur children with dyslexia, as well as to research into dyslexia and different racial genetics. Moreover, to better explicate the role of KIAA0319 underlying dyslexia etiology and pathology, more functional studies in Uyghur population need to be conducted.

## ACKNOWLEDGMENTS

This study was supported in part by National Natural Science Foundation of China(81360434). We wish to thank Dr. Da Ding and Dr. Yan Liu (Genesky Biotechnologies Inc., Shanghai, China) for technical support.

